# A robust and efficient method for Mendelian randomization with hundreds of genetic variants: unravelling mechanisms linking HDL-cholesterol and coronary heart disease

**DOI:** 10.1101/566851

**Authors:** Stephen Burgess, Christopher N Foley, Elias Allara, James R Staley, Joanna MM Howson

## Abstract

Mendelian randomization (MR) investigations with large numbers of genetic variants are becoming increasingly common. However, the reliability of findings from a MR investigation is dependent on the validity of the genetic variants as instrumental variables. We developed a method to identify groups of genetic variants with similar causal effect estimates, which may represent distinct mechanisms by which the risk factor influences the outcome. Our contamination mixture method is a robust and efficient method for valid MR in the presence of invalid IVs. Compared to other robust methods, our method had the lowest mean squared error across a range of realistic scenarios. The method is fast and efficient, and can perform analysis with hundreds of variants in a fraction of a second. In a MR analysis for high-density lipoprotein (HDL) cholesterol and coronary heart disease (CHD) risk, the method identified 11 variants associated with increased HDL-cholesterol, decreased triglyceride levels, and decreased CHD risk that had the same directions of associations with platelet distribution width and other blood cell traits, suggesting a shared mechanism linking lipids and CHD risk relating to platelet aggregation.

## Introduction

Distinguishing between correlation and causation is a fundamentally important problem when trying to understand disease mechanisms. Mendelian randomization is an approach to assess whether a risk factor has a causal effect on an outcome based on observational data [1, 2]. Mendelian randomization can exploit summarized data on genetic associations obtained from genome-wide association studies to link modifiable risk factors to disease outcomes [3, 4]. However, the validity of findings from a Mendelian randomization investigation is not guaranteed.

Mendelian randomization relies on genetic variants satisfying the assumptions of an instrumental variable (IV) [5]. A valid IV must be associated with the risk factor of interest in a specific way, such that it is not also associated with confounders of the risk factor–outcome association, nor does it affect the outcome directly (but only potentially indirectly via its effect on the risk factor of interest). An IV can be used to estimate the average causal effect of the risk factor on the outcome [6]. If there are multiple valid IVs, then the estimates from each IV should be similar to each other [7]. If estimates differ substantially, then it is likely that not all the IVs are valid.

Genetic variants represent a fertile source of candidate IVs, particularly for genes that have well-understood functions and specific effects on risk factors [8]. Genetic variants are fixed at conception, providing some immunity to reverse causation (as the genetic variant must temporally precede the outcome) and confounding (as the genetic code cannot be influenced by confounding factors that act after conception). However, there are also several reasons why genetic variants may not be valid IVs: such as linkage disequilibrium with another functional variant, pleiotropy (that is, a variant affects risk factors on different causal pathways), and population stratification [9, 10, 11].

Several approaches for making inferences with some invalid instruments have previously been proposed. These include methods that assume that a majority of the candidate instruments are valid IVs [12, 13, 14], and those that assume a plurality of the candidate instruments are valid IVs [15, 16]. The plurality assumption means that out of all groups of candidate instruments having the same asymptotic causal estimate, the largest group is the group of valid IVs. A similar assumption is made in outlier-removal methods, such as MR-PRESSO, which sequentially removes candidate instruments from the analysis based on a heterogeneity measure until all the remaining variants have similar estimates [17]. Other assumptions have also been made: for example, the MR-Egger method assumes that the distribution of direct effects of candidate instruments on the outcome is independent from the distribution of associations with the risk factor (referred to as the Instrument Strength Independent of Direct Effect – InSIDE – assumption) [18].

We here introduce the contamination mixture method as a method for obtaining valid causal inferences with some invalid IVs. Compared to other approaches for robust instrumental variable analysis, we believe our proposal has a number of advantages in giving asymptotically consistent estimates under the ‘plurality of valid instruments’ assumption, being fully likelihood-based, being computationally scalable to large numbers of candidate instruments, and being implemented using summarized genetic data that are widely available from large consortia. A particular feature of the proposed method is that it can identify groups of genetic variants with similar causal estimates. If multiple such groups are identified, this suggests that there may be several causal mechanisms associated with the same risk factor that affect the outcome.

In this paper, we compare the performance of the proposed contamination mixture method in an extensive simulation study with realistic parameters. We show that our new method performs well compared to previously proposed methods in terms of bias, Type 1 error rate, and efficiency. We then illustrate the use of the method in an example considering the causal effect of high-density lipoprotein (HDL) cholesterol on coronary heart disease (CHD) risk, demonstrating a bimodal distribution of the variant-specific estimates. We investigate factors that identify a group of genetic variants associated with HDL-cholesterol and triglyceride levels having a strong protective effect on CHD risk, showing that several of these variants have the same directions of associations with various blood cell traits. We then discuss the implications for identifying causal risk factors and mechanisms.

## Results

### Overview of the proposed contamination mixture method

The contamination mixture method is implemented by constructing a likelihood function based on the variant-specific causal estimates. If a genetic variant is a valid instrument, then its causal estimate will be normally distributed about the true value of the causal effect. If a genetic variant is not a valid instrument, then its causal estimate will be normally distributed about some other value. We assume that the values estimated by invalid instruments are normally distributed about zero with a large standard deviation. This enables a likelihood function to be specified that is a product of two-component mixture distributions, with one mixture distribution for each variant. The computational time for maximizing this likelihood directly is exponential in the number of genetic variants. We use a profile likelihood approach to reduce the computational complexity to be linear in the number of variants (Methods).

Briefly, we consider different values of the causal effect in turn. For each value, we calculate the contribution to the likelihood for each genetic variant as a valid instrument and as an invalid instrument. If the contribution to the likelihood as a valid instrument is greater, then we take the variant’s contribution as a valid instrument; if less, then its contribution is taken as an invalid instrument. This gives us the configuration of valid and invalid instruments that maximizes the likelihood for the given value of the causal effect. This is a profile likelihood, a one-dimensional function of the causal effect. The point estimate is then taken as the value of the causal effect that maximizes the profile likelihood. A 95% confidence interval is constructed by taking the set of values of the causal effect for which twice the difference between the log-likelihood calculated at the value and at the maximum is less than the 95th percentile of a chi-squared distribution with one degree of freedom. We note that the confidence interval from this approach is not constrained to be symmetric or even a single range of values. A confidence interval consisting of multiple disjoint ranges would occur if there were multiple groups of genetic variants having estimates that are mutually consistent within the group, but different between the groups.

### Comparison with previous methods

To compare the performance of the contamination mixture method against other robust methods for Mendelian randomization, we performed a simulation study with a broad range of realistic scenarios. We simulated data in a two-sample setting under 4 scenarios: 1) all genetic variants are valid IVs, 2) invalid IVs have balanced pleiotropic direct effects on the outcome, 3) invalid IVs have directional pleiotropic direct effects on the outcome, and 4) invalid IVs have directional pleiotropic effects on the outcome via a confounder. In the first three scenarios, the InSIDE assumption is satisfied, while in the fourth it is not. We took 100 genetic variants as candidate instruments, with the number of invalid IVs in Scenarios 2 to 4 being 20, 40 or 60. We performed several methods: the standard inverse-variance weighted (IVW) method that assumes all genetic variants are valid IVs [19], a weighted median method that assumes that a majority of genetic variants are valid IVs [14], the MR-Egger method [18], a weighted mode-based estimation method that assumes a plurality of genetic variants are valid IVs [16], MR-PRESSO [17], and the proposed contamination mixture method. 10 000 simulated datasets were analysed for each scenario (Methods).

Table 1 shows that when all variants are valid IVs, all methods give unbiased estimates. The most efficient robust method, judged by standard deviation of the causal estimates and empirical power to detect a true effect, is the MR-PRESSO method. This method is similar in efficiency to the IVW method, which is optimally-efficient with all valid IVs [20], but biased when one or more genetic variants are invalid IVs. The contamination mixture and weighted median methods are slightly less efficient, while the mode-based method and MR-Egger methods are considerably less efficient. Mean estimates are attenuated towards the null in all methods due to weak instrument bias [21]; attenuation was more severe in the MR-Egger and mode-based methods (Table 1).

**Table 1:** Comparison of methods when all genetic variants are valid instruments: All methods tested gave unbiased causal estimates in the simulation scenario when there were no invalid instruments. Mean, standard deviation (SD), mean standard error (mean SE) of estimates, and empirical power (%) in simulation study for Scenario 1 (all 100 variants valid instruments). Abbreviation: MR-PRESSO = Mendelian Randomization Pleiotropy RESidual Sum and Outlier method of Verbanck et al. [17].

| Method | Scenario 1: all instruments valid |  |  |  |
| --- | --- | --- | --- | --- |
|  | Mean | SD | Mean SE | Power |
| Null causal effect: $\theta = 0$ | | | | |
| Inverse-variance weighted | 0.000 | 0.022 | 0.023 | 4.2 |
| MR-Egger | 0.001 | 0.061 | 0.062 | 4.4 |
| Weighted median | 0.000 | 0.028 | 0.033 | 2.1 |
| MR-PRESSO | -0.000 | 0.022 | 0.022 | 5.1 |
| Weighted mode-based | 0.001 | 0.061 | 0.199 | 0.3 |
| Contamination mixture | 0.000 | 0.028 | - | 4.9 |
| Positive causal effect: $\theta = +0.1$ | | | | |
| Inverse-variance weighted | 0.094 | 0.023 | 0.024 | 98.1 |
| MR-Egger | 0.063 | 0.064 | 0.066 | 16.1 |
| Weighted median | 0.091 | 0.030 | 0.035 | 78.8 |
| MR-PRESSO | 0.094 | 0.024 | 0.023 | 98.0 |
| Weighted mode-based estimate | 0.082 | 0.051 | 0.143 | 15.5 |
| Contamination mixture | 0.096 | 0.029 | - | 91.0 |

Table 2 shows that no one method outperforms others in every scenario. The MR-Egger method performs well in terms of bias under the null and Type 1 error rate in Scenarios 2 and 3, but has the lowest power to detect a positive effect in these scenarios, and is the most biased method in Scenario 4. The weighted median method has lower bias than the IVW method and reasonable power to detect a causal effect, but has high Type 1 error rate in Scenarios 3 and 4 even with only 20 invalid instruments. The MR-PRESSO method is the most efficient at detecting a causal effect, but also has high Type 1 error rate in Scenarios 3 and 4 even with only 20 invalid instruments. The weighted mode-based estimation method generally has low bias and low Type 1 error rate inflation with up to 40 invalid instruments, but also has low power to detect a causal effect. The contamination mixture method generally has good properties, with low bias and low Type 1 error rate inflation for up to 40 invalid instruments, but much better power than the mode-based estimation method to detect a causal effect. Performance with 60 invalid instruments is generally poor for all methods.

**Table 2:**
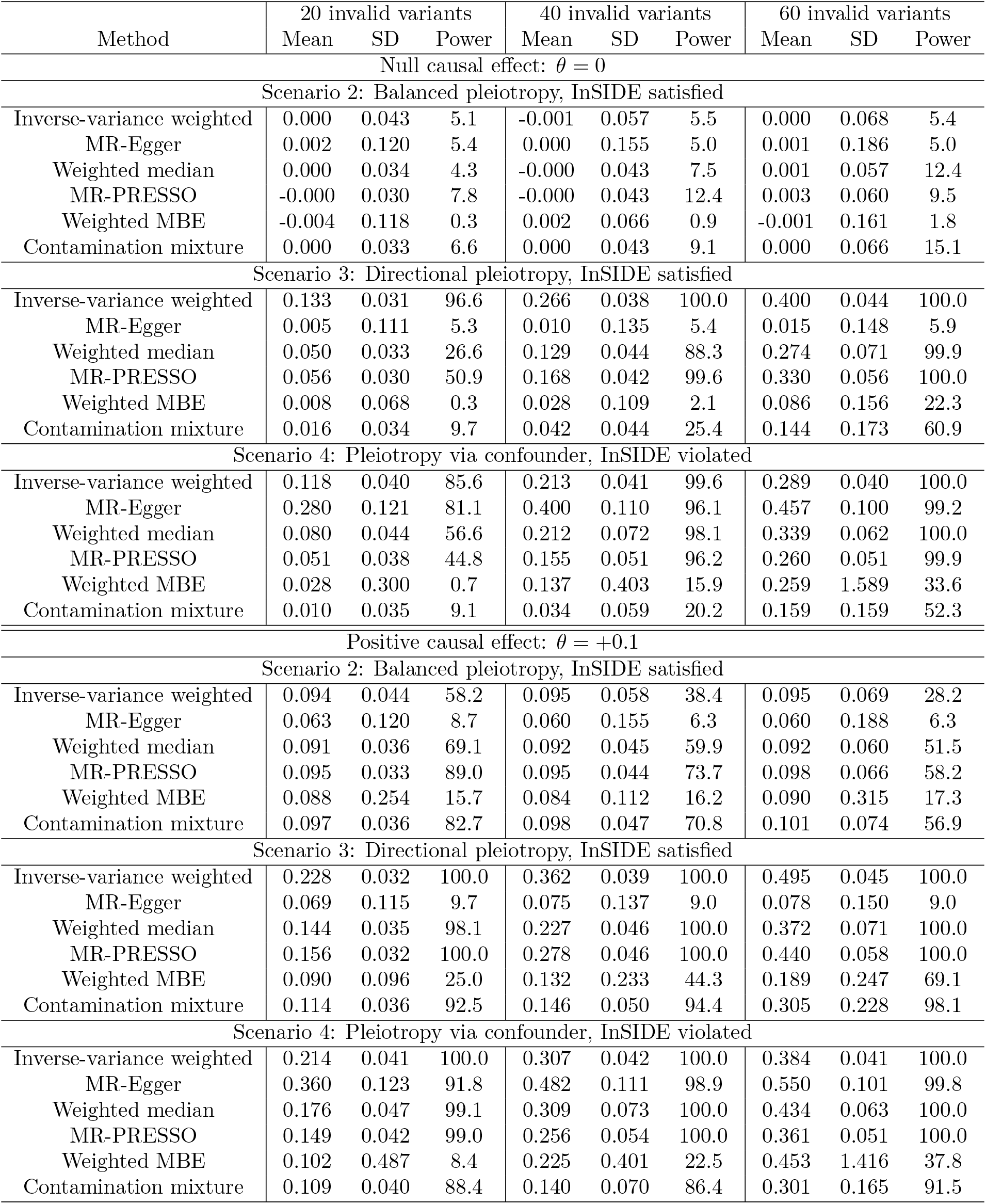
Comparison of methods with some invalid instruments: No one method outperformed all others in every scenario, but the contamination mixture method had good overall performance across scenarios with up to 40 invalid instruments out of 100. Performance with 60 invalid instruments was generally poor for all methods. Mean, standard deviation (SD) of estimates, and empirical power (%) in simulation study for Scenarios 2, 3, and 4. Abbreviations: MR-PRESSO = Mendelian Randomization Pleiotropy RESidual Sum and Outlier method of Verbanck et al. [17]; MBE = mode-based estimate of Hartwig et al. [16].

The mean squared error of each method in scenarios with a null causal effect is shown in Figure 1. The contamination mixture method clearly dominates other methods in terms of overall performance according to this measure, particularly in scenarios with 40 or 60 invalid instruments.

**Figure 1:**
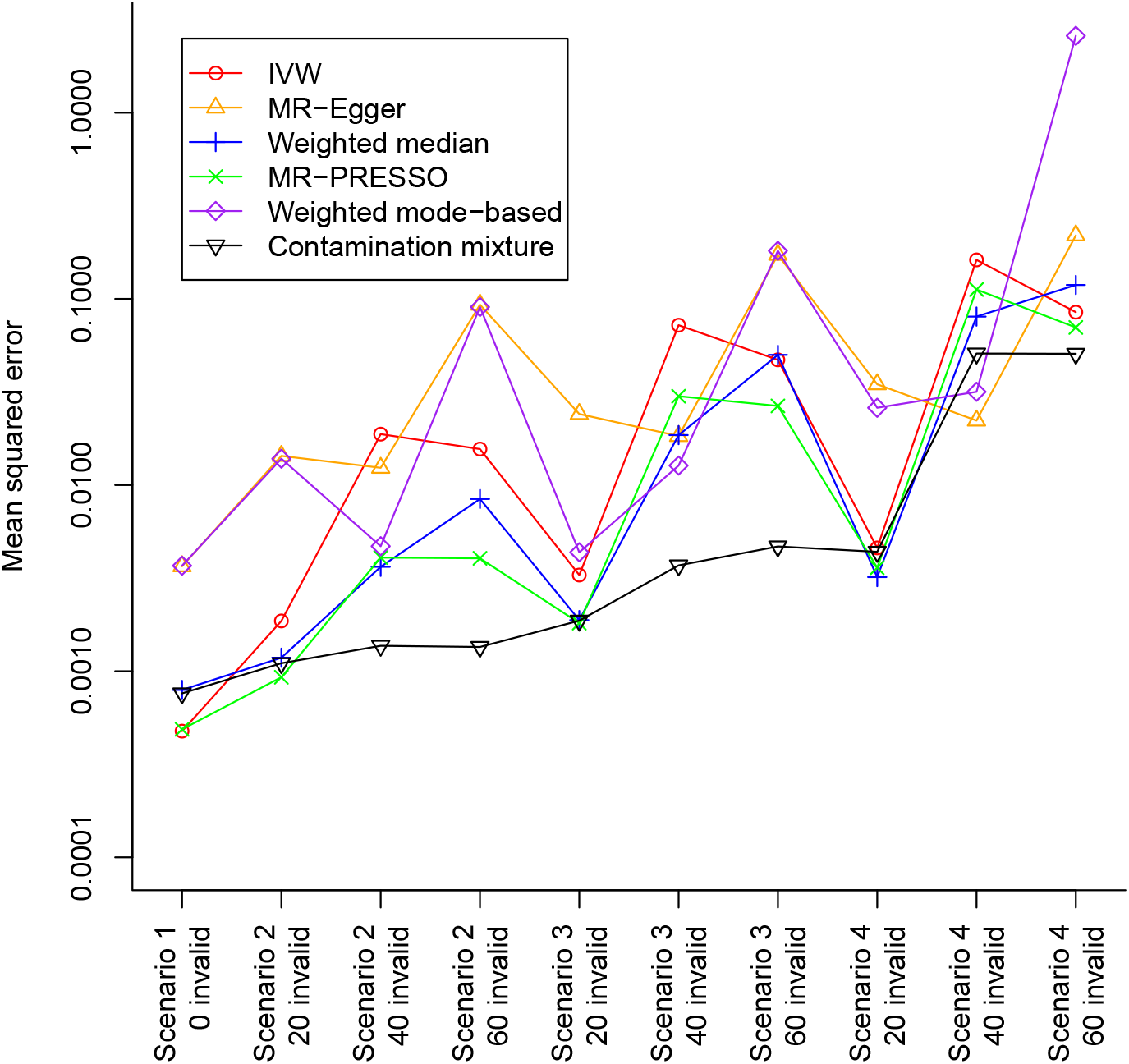
Comparison of the methods based on mean squared error criterion: The mean squared error of the various methods is plotted in each scenario with a null causal effect. The corresponding plot with a positive causal effect was practically identical. The contamination mixture method has the best overall performance according to this measure, particularly in scenarios where 40 or 60 out of the 100 genetic variants were invalid instruments. The vertical axis is plotted on a logarithmic scale.

### Unravelling the effect of HDL-cholesterol on coronary heart disease risk

To consider the causal effect of HDL-cholesterol on CHD risk, we took 86 uncorrelated genetic variants previously associated with HDL-cholesterol at a genome-wide level of significance in the Global Lipids Genetic Consortium (GLGC) [22]. Associations with HDL-cholesterol were estimated in the GLGC based on up to 188,577 individuals of European ancestry, and associations with CHD risk from the CARDIoGRAMplusC4D consortium on up to 60,801 CHD cases and 123,504 controls predominantly of European ancestry [23] (Supplementary Table A1). Previous analyses for HDL-cholesterol with these variants using the IVW method indicated a protective causal effect, whereas analyses using robust methods (in particular, weighted median and MR-Egger) suggest that the true effect is null [24]. A null effect has been observed in most trials for CETP inhibitors that raise HDL-cholesterol [25, 26]. In one trial a modest protective effect was observed [27], although this may be ascribed to the LDL-cholesterol lowering effect of the drug.

The contamination mixture method gives a likelihood function that is bimodal (Supplementary Figure A1), with a point estimate (representing the odds ratio for CHD per 1 standard deviation increase in HDL-cholesterol) of 0.67 for the primary maximum and 0.93 for the secondary maximum and a 95% confidence interval comprising two disjoint ranges from 0.59 to 0.77, and from 0.88 to 0.96. Visual inspection of the scatter graph (Figure 2) reveals there are several variants suggesting a protective effect of HDL-cholesterol on CHD risk. The two regions of the confidence interval for the causal effect suggests that the variants can be divided into at least two groups, and the method is uncertain which of the groups is larger. However, 43 of the 86 genetic variants are also associated with triglycerides at *p* < 10^−5^, meaning that the associations with CHD risk may be driven by a harmful effect of triglycerides rather than a protective effect of HDL-cholesterol, as suggested in multivariable Mendelian randomization analyses [28]. Still, it is worthwhile investigating those variants that evidence the strong protective effect, to see if there is any commonality between them that may suggest a causal mechanism.

**Figure 2:**
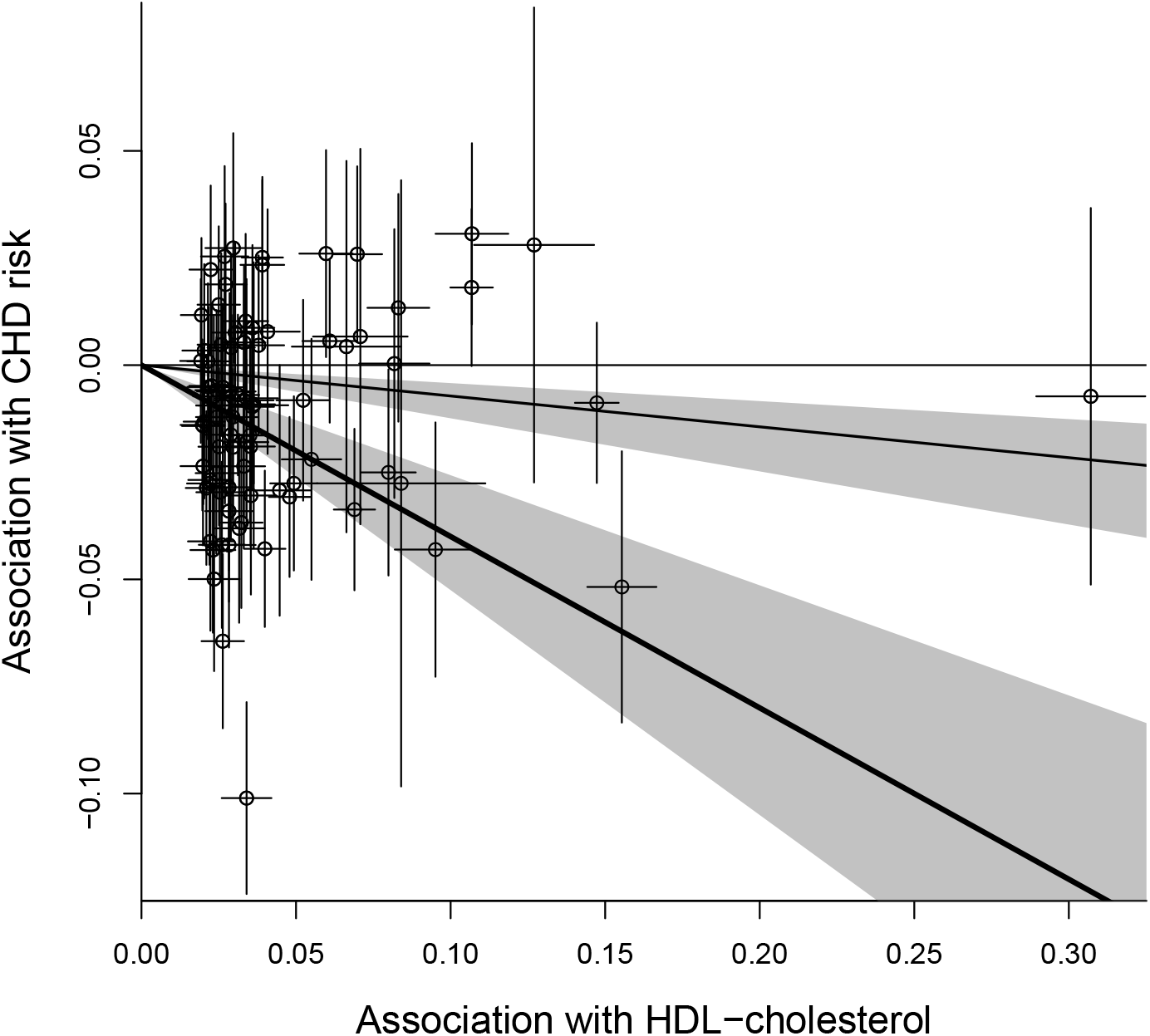
Scatter plot of genetic associations: Genetic associations with HDL-cholesterol (standard deviation units) against genetic associations with CHD risk (log odds ratios). Error bars for genetic associations are 95% confidence intervals. Heavy black line is the causal estimate from the contamination mixture method with the strongest signal, lighter black line is the causal estimate from the secondary peak. The grey area is the 95% confidence interval for the causal effect; this comprises two ranges as the likelihood is bimodal.

We searched for associations of all 86 variants in PhenoScanner [29], a database of summarized data from genome-wide association studies, and found 3209 traits for which at least one genetic association was available. for each variant, we calculated the posterior probability of being a valid genetic variant given a causal effect of 0.67, the estimate from the contamination mixture method. Traits were then ranked according to the mean posterior probability for all variants associated with the trait (Methods, Supplementary Figure A2). The top ranked trait was platelet distribution width. The next ranked traits were also blood cell traits: mean corpuscular hemoglobin concentration, and red cell distribution width (Supplementary Table A2). This suggests that the potential mechanism linking these variants to CHD risk relates to blood cell trait-related mechanisms.

We investigated variants that were associated with increased HDL-cholesterol, decreased CHD risk, and at least one of the above blood cell traits (Supplementary Figure A3). We found 11 genetic variants in 9 distinct gene regions (including those having the largest posterior probability) with a distinct pattern of associations: decreased triglycerides, decreased mean corpuscular hemoglobin concentration, decreased platelet distribution width, and increased red cell distribution width (Table 3). The similarity in the presence and direction of associations with these traits suggests a potential shared causal mechanism. In particular, the finding of platelet distribution width suggests a link with platelet aggregation, a known risk factor for CHD that has previously been linked to HDL-cholesterol [30]. To further investigate evidence of a shared causal mechanism, we performed multi-trait colocalization across these traits for each of the gene regions. For 7 of the regions, there was strong evidence of colocalization for HDL-cholesterol, CAD risk, and at least one of the blood cell traits (Supplementary Table A3).

**Table 3:**
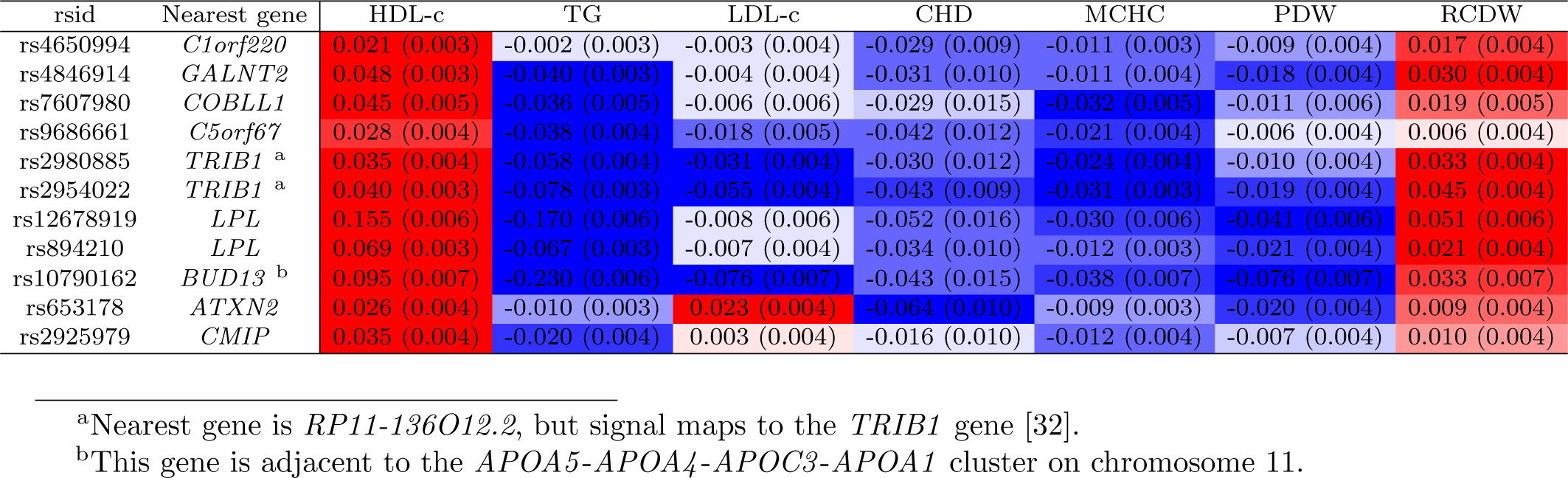
Variants having same directions of associations with HDL-cholesterol, coronary heart disease, and blood cell traits: Details of genetic variants, nearest gene, beta-coefficients (standard errors, SE) for associations with HDL-cholesterol (HDL-c), triglycerides (TG), LDL-cholesterol (LDL-c), coronary heart disease (CHD) risk, mean corpuscular hemoglobin concentration (MCHC), platelet distribution width (PDW), and red cell distribution width (RCDW) for 11 genetic variants in 9 distinct gene regions having a distinct pattern of associations. All associations are orientated to the HDL-cholesterol increasing allele. Red indicates that the association is positive; blue for negative. The brightness of the coloring corresponds to the p-value for the strength of association; brighter colors correspond to lower p-values.

## Discussion

In this paper, we have introduced a new robust method for Mendelian randomization analysis referred to as the contamination mixture method. Compared to other robust methods, the contamination mixture method had the best all-round performance in a simulation study – maintaining little bias and close to nominal Type 1 error rates under the null with up to 40 invalid genetic variants out of 100, having reasonable power to detect a causal effect in all scenarios, and having the lowest average mean squared error of all methods. In a Mendelian randomization analysis for HDL-cholesterol on CHD risk, the method detected two separate groups of variants suggesting distinct protective effects on CHD risk. A hypothesis-free search of traits revealed that several variants in the group with the stronger protective effect were associated with platelet distribution width and two other blood cell traits, with consistent directions of association across 11 variants in 9 gene regions. This suggests that the apparent protective effect of these variants may be driven by a shared mechanism, potentially relating to platelet aggregation. Overall, the proposed method is able to make reliable causal inferences in Mendelian randomization investigations with some invalid instruments, as well as highlighting when there are multiple causal effects represented in the data that may be driven by different mechanisms.

Rather than attempting to model the distribution of the estimates from invalid genetic variants, the contamination mixture method proposes that these estimates come from a normal distribution centred at the origin with a wide standard deviation that is pre-specified by the user. We experimented with alternative methods that estimate this distribution. However, as the identity of the invalid instruments is unknown, the performance of these methods was worse than the approach presented here. The runtime of our proposed method is low – on an Intel i7 2.70 GHz processor, the contamination mixture method took 0.08 seconds to analyse one of the simulated datasets with 100 variants. In comparison, the MR-PRESSO method took 120 seconds, and the mode-based estimation method 81 seconds. The complexity of the contamination mixture model is linear in the number of genetic variants, so the benefit in computational time over other methods would be even greater if more genetic variants were included in the analysis. The previously proposed heterogeneity-penalized method [31], on which the contamination mixture method is based, would be prohibitively slow even with 30 genetic variants. A disadvantage of our proposed method is sensitivity to the standard deviation parameter. In the example of HDL-cholesterol and CHD risk, while the two distinct groups of genetic variants were detected at some values of the standard deviation parameter, at other values distinct groups were not detected.

As genome-wide association studies become larger, the number of genetic variants associated with different traits increases, and it is increasingly unlikely that all these genetic variants are valid instruments for the risk factor of interest. For risk factors with dozens or even hundreds of associated variants, a paradigm shift is required away from approaches that assume the majority of genetic variants are valid instruments, and towards methods that attempt to find genetic variants with similar estimates that might represent a particular causal mechanism. Heterogeneity in variant-specific estimates is inevitable, but heterogeneity should be seen as an opportunity to find causal mechanisms, rather than a barrier to Mendelian randomization investigations. In our example, we demonstrated how a group of genetic variants having similar causal effects for HDL-cholesterol on CHD risk are linked by their association with platelet distribution width, suggesting a possible mechanism linking lipids to CHD risk related to platelet aggregation. While further research is needed to establish this mechanism, the approach demonstrates the possibility of finding information on causal mechanisms amongst heterogeneous data.

In conclusion, we have introduced a robust method for Mendelian randomization that outperforms other methods (including MR-PRESSO) in terms of all-round performance, and can identify groups of variants having similar causal estimates, enabling the identification of causal mechanisms.

### Software

Code for the contamination mixture method is presented in the Supplementary Material and is implemented in the MendelianRandomization package (https://cran.r-project.org/web/packages/MendelianRandomization/index.html).

## Online Methods

### Instrumental variable assumptions

A genetic variant is an instrumental variable if:

1. it is associated with the risk factor of interest,
2. it is not associated with any confounder of the risk factor–outcome association, and
3. it is not associated with the outcome except potentially indirectly via the risk factor [8].

We use the term ‘candidate instrumental variable’ to indicate a variable that is treated as an instrumental variable, without prejudicing whether it satisfies the instrumental variable assumptions or not.

We assume that the associations between the instrumental variable and risk factor, and the instrumental variable and outcome are linear and homogeneous between individuals with no effect modification, and the causal effect of the risk factor on the outcome is linear [33]. These assumptions are not necessary for the instrumental variable estimate to be a valid test of the causal null hypothesis, but they ensure that all valid instrumental variables estimate the same causal effect.

If summarized data are available representing the association of genetic variant *j* with the risk factor (beta-coefficient 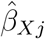 and standard error 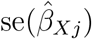), and the association with the outcome (beta-coefficient 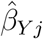) and standard error 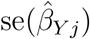, then the causal effect of the risk factor on the outcome 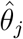 can be estimated as:

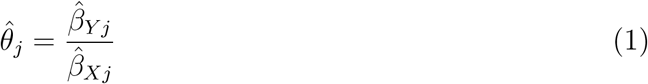

and its standard error 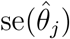 as:

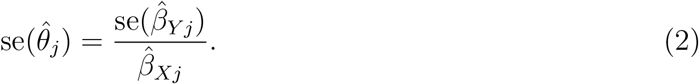

We refer to 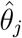 as a variant-specific causal estimate. The formula for the standard error only takes into account the uncertainty in the genetic association with the outcome, but this is typically much greater than the uncertainty in the genetic association with the risk factor, and does not tend to lead to Type 1 error inflation [19]. (In fact, accounting for this uncertainty naively leads to a correlation between the estimated association with the risk factor and the standard error of the variant-specific estimate – which can have more serious consequences in practice than ignoring this source of uncertainty [34, 35].)

### Estimation with multiple instrumental variables

If there are multiple instrumental variables, then a more precise estimate of the causal effect can be obtained using information on all the instrumental variables. If individual-level data are available, the two-stage least squares method is performed by regressing the risk factor on the instrumental variables, and then regressing the outcome on fitted values of the risk factor from the first stage regression. The same estimate can be obtained by taking a weighted mean of the variant-specific causal estimates using inverse variance weights, as in a meta-analysis [36]. The inverse variance weighted (IVW) estimate can be expressed as:

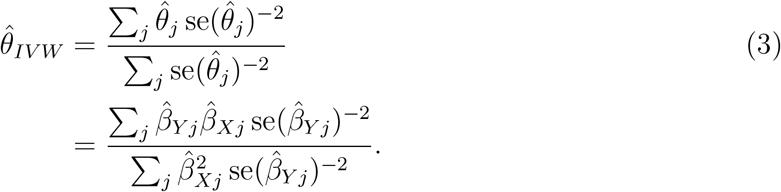

The IVW estimate can also be obtained by weighted regression using the following model:

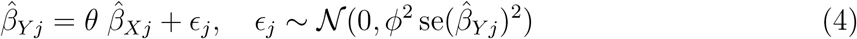

Here, we include an additional term *ϕ*, which is the residual standard error in the regression model. In a fixed-effect analysis, this parameter is fixed to be one, but in a random-effects analysis, we allow this term (which represents overdispersion of the variance-specific causal estimates) to be estimated (although we do not allow it to take values less than one, as underdispersion is implausible) [37, 38].

The two-stage least squares method (and hence the fixed-effect IVW method) is the most efficient unbiased combination of the variant-specific estimates [20]. However, it is only a consistent estimate of the causal effect if all the candidate instrumental variables are valid. We introduce the contamination mixture method as a robust method for instrumental variable analysis (a robust method is a method that provides asymptotically consistent estimates under weaker assumptions that all genetic variants being valid instruments).

### Contamination mixture method

Suppose we have *J* genetic variants that are candidate instrumental variables. If genetic variant *j* is a valid instrumental variable, then the variant-specific estimate from that variant 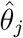 is normally distributed about the true causal parameter *θ* with standard deviation equal to the standard error of the ratio estimate 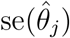:

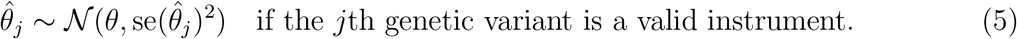

If the same variant is not a valid instrumental variable, then the ratio estimate from that variant 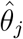 is normally distributed about some other value *θ_F,j_* with standard deviation equal to the standard error of the ratio estimate 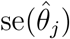. We assume that these *θ_F,j_* are normally distributed about zero with standard deviation *ψ*, such that the distribution of 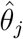 is:

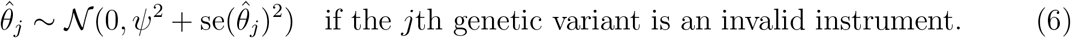

We assume that the standard errors 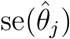 are fixed and known. Our method makes no attempt to model the distribution of the estimates from invalid instruments. We simply propose a symmetric normal distribution of the estimates about the origin, with the variance of the distribution comprising the proposed variability in the estimands *θ_F,j_* (which is *ψ*^2^) plus the uncertainty in the estimate about its asymptotic value (which is the square of its standard error). If the variant-specific estimate is close to the proposed causal parameter *θ*, then the likelihood corresponding to the genetic variant being a valid instrument will be larger; if the variant-specific estimate is not close to the proposed causal parameter *θ*, then the likelihood corresponding to the genetic variant being a invalid instrument will be larger.

We notate the model (that is, the configuration of valid and invalid instruments) as a vector *γ*, where *γ_j_* = 1 when genetic variant *j* is valid, and *γ_j_* = 0 otherwise. The likelihood function is then:

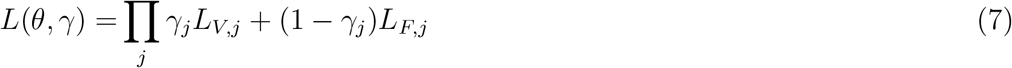

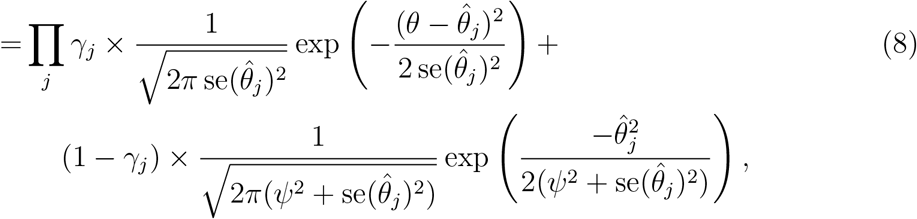

where the likelihood contribution from each genetic variant is *L_V,j_* if genetic variant *j* is valid and *L_F,j_* if genetic variant *j* is invalid. Making inferences using this likelihood is not simple, as the parameter space for *γ* grows exponentially with the number of genetic variants, and there is no guarantee that the likelihood will be unimodal, making it difficult to apply stochastic approaches to explore the model space.

We proceed using a profile likelihood approach. If the causal estimate *θ* is fixed, then the optimal model *γ* to maximize the likelihood is clear: we should take *γ_j_* = 1 if *L_V,j_* > *L_F,j_* and *γ_j_* = 0 otherwise. We denote this model choice as 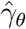. This implies we can easily calculate a profile likelihood 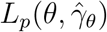 for any value of *θ*. We perform inferences on *θ* by calculating this profile likelihood at a range of values of *θ*. The causal estimate 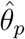 is taken as the value of *θ* that maximizes the profile likelihood, and the confidence interval as the values of *θ* such that:

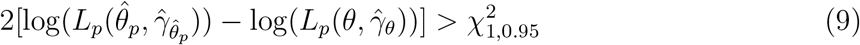

where 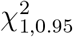 is the 95th percentile of a chi-squared distribution with 1 degree of freedom. The confidence interval is based on Wilks’ likelihood ratio test, and is not constrained to be symmetric or a single range of values. We note that the profile likelihood is a continuous function of *θ*: it is clearly a continuous function for ranges of *θ* when 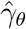 does not vary, and elements *γ_j_* only change their value when the profile likelihood is the same for *γ_j_* = 0 and *γ_j_* = 1.

To allow for excess heterogeneity in estimates of the true causal parameter (analogous to a random-effects model for the IVW method), we estimate an overdispersion parameter as the residual standard error *ϕ* from weighted regression of the genetic associations with the outcome on the genetic associations with the exposure using the genetic variants judged to be valid at the causal estimate 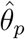 as in the inverse-variance weighted method. We then replace 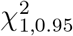 in the above formula with 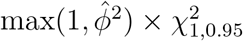.

We recommend taking the value of *ψ* to be the standard deviation of the ratio estimates based on all the genetic variants, multiplied by 1.5. This value was taken in the applied example. This means that the standard deviation of invalid estimands *θ_F,j_*, is guided by the variability of the observed ratio estimates, but inflated as the valid instruments will have more similar causal estimates. However, a sensitivity analysis for this parameter is advised. In the applied example, the causal estimate, as well as whether the method detected two separate groups of variants or not, was sensitive to the choice of this parameter (Supplementary Table A4).

### Simulation study

To compare the contamination mixture method with previously developed methods for Mendelian randomization, we perform a simulation study. We consider four scenarios:

1. no pleiotropy – all genetic variants are valid instruments;
2. balanced pleiotropy – some genetic variants have direct (pleiotropic) effects on the outcome, and these pleiotropic effects are equally likely to be positive as negative;
3. directional pleiotropy – some genetic variants have direct (pleiotropic) effects on the outcome, and these pleiotropic effects are simulated to be positive;
4. pleiotropy via a confounder – some genetic variants have pleiotropic effects on the outcome via a confounder. These pleiotropic effects are correlated with the instrument strength.

In the first three scenarios, the Instrument Strength Independent of Direct Effect (InSIDE) assumption [18] is satisfied; in Scenario 4, it is violated. This is the assumption required for the MR-Egger method to provide consistent estimates.

We simulate data for a risk factor *X*, outcome *Y*, confounder *U* (assumed unmeasured), and *J* genetic variants *G_j_, j* = 1,…, *J*. Individuals are indexed by *i*. The data-generating model for the simulation study is as follows:

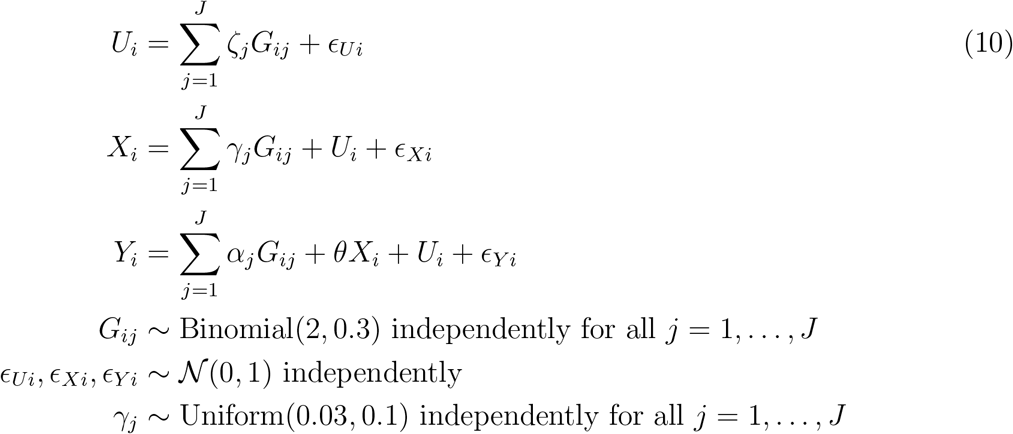

The risk factor and outcome are positively correlated due to confounding even when the causal effect *θ* is zero through the unmeasured confounder *U*. The genetic variants are modelled as single nucleotide polymorphisms (SNPs) with a minor allele frequency of 30%. A total of *J* = 100 genetic variants are used in each analysis. For each of Scenarios 2 to 4, we considered cases with 20, 40 and 60 invalid instruments. For valid instruments, the *α_j_* and *ζ_j_* parameters were set to zero. For invalid instruments, the *α_j_* parameters were either drawn from a uniform distribution on the interval from –0.1 to 0.1 (Scenario 2), or from 0 to 0.1 (Scenario 3), or set to zero (Scenario 4). The *ζ_j_* parameters were either set to zero (Scenarios 2 and 3), or drawn from a uniform distribution on the interval from –0.1 to 0.1 (Scenario 4). The causal effect θ was either set to 0 (no causal effect) or +0.1 (positive causal effect). The *γ_j_* parameters were drawn from a uniform distribution on 0.03 to 0.1, meaning that the average value of the *R*^2^ statistic for the 100 variants across simulated datasets was 9.3% (from 10.5 to 12.7% in Scenario 4) corresponding to an average F statistic of 20.5 (from 23.3 to 28.9 in Scenario 4).

In total, 10 000 datasets were generated in each scenario. We considered a two-sample setting in which genetic associations with the risk factor and outcome were estimated on non-overlapping groups of 20 000 individuals. We compared estimates from the proposed contamination mixture method with those from a variety of methods: the standard IVW method, MR-Egger [18] (both using random-effects), the weighted median method [14], MR-PRESSO [17], and the mode-based estimation (MBE) method of Hartwig et al. [16]. To avoid extreme values in a minority of analyses, we set *ψ* = 1 in the contamination mixture method in all analyses. Each of the methods was implemented using summarized data only. Default values were used by the methods; in particular, the MBE method was implemented using the weighted option with bandwidth *ϕ* =1 under the no measurement error (NOME) assumption (for similarity and thus comparability with other methods), and the MR-PRESSO method was performed trimming variants at a p-value threshold of 0.05.

For computational reasons (the methods took over 100 times longer to run than the other methods put together), the MR-PRESSO and MBE methods were performed on 1000 datasets per scenario only. Mean squared error was calculated in each scenario by averaging across the 10 000 datasets (1000 datasets for the MR-PRESSO and MBE methods).

### Calculation of the posterior probability for HDL-cholesterol associated variants

The posterior probability of a genetic variant being a valid instrumental variable can be calculated as:

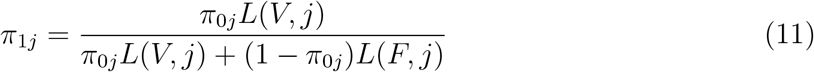

where *π*_0*j*_ is the prior probability of being a valid instrument. We take *π*_0*j*_ to be the absolute value of the association of variant j with HDL-cholesterol in standard deviation units. This ensures that variants having greater association with the risk factor receive a higher prior weight. If the prior weights for all variants were equal, then a variant having a weak association with the risk factor and an imprecise association with the outcome that is compatible with multiple values of the causal effect could receive the same posterior weight as a variant having a strong association with the risk factor and a precise association with the outcome that is compatible with a far narrow range of values of the causal effect.

### Colocalization for HDL-cholesterol associated variants

Multi-trait colocalization was performed using the Hypothesis Prioritization Colocalization (HyPrColoc) package (https://github.com/jrs95/hyprcoloc). This package performs multitrait colocalization in a similar way to *moloc*, the multi-trait extension to *coloc* [39], but in a computationally efficient way that allows colocalization of large numbers of traits to be performed. We investigated colocalization between six traits: HDL-cholesterol, triglycerides, CHD risk, mean corpuscular hemoglobin concentration, platelet distribution width, and red cell distribution width. These blood cell traits were selected as variants associated with these traits have the greatest mean posterior probability of belonging to the largest group of variants identified by the contamination mixture method (Supplementary Table A2). Associations with the blood cell traits were estimated in 173,480 unrelated European-descent individuals from the UK Biobank and INTERVAL studies [40]. For each gene region, we took all available variants from the relevant recombination window around the gene [41]. Colocalization was performed using default settings for the priors in the *hyprcoloc* function (prior probability of initial trait association 0.0001, conditional probability of subsequent trait having shared association 0.02), and with the uniform priors setting as the default setting can be overly conservative.

While the exact pattern of colocalization differed between the gene regions, colocalization between HDL-cholesterol, CHD risk, and at least one blood cell trait was observed for 3 of the gene regions using the conservative priors, and for 7 regions using uniform priors (Supplementary Table A3). The posterior probability of colocalization was at least 0.7 in all cases, except when using conservative priors in the *C5orf67* gene region. For this region, there was evidence of colocalization between HDL-cholesterol, triglycerides, CHD risk, and mean corpuscular hemoglobin concentration at posterior probability 0.59, and evidence of colocalization between HDL-cholesterol, triglycerides, and mean corpuscular hemoglobin concentration only (excluding CHD risk) at posterior probability 0.96. For the two gene regions that did not show evidence of colocalization between these traits, one possible explanation is the presence of multiple causal variants in the region; as the *ATXN2* gene region reported colocalization between HDL-cholesterol and CHD risk, and separately between the blood cell traits. For *COBLL1*, there was colocalization between HDL-cholesterol and the blood cell traits, but not CHD risk.

## Acknowledgements

This work was funded by the UK Medical Research Council (MR/L003120/1, MC_UU_00002/7), British Heart Foundation (RG/13/13/30194), and the UK National Institute for Health Research Cambridge Biomedical Research Centre. Stephen Burgess is supported by Sir Henry Dale Fellowship jointly funded by the Wellcome Trust and the Royal Society (Grant Number 204623/Z/16/Z). The views expressed are those of the authors and not necessarily those of the National Health Service, the National Institute for Health Research or the Department of Health and Social Care.

## Supplementary Material

### A.1 Software code

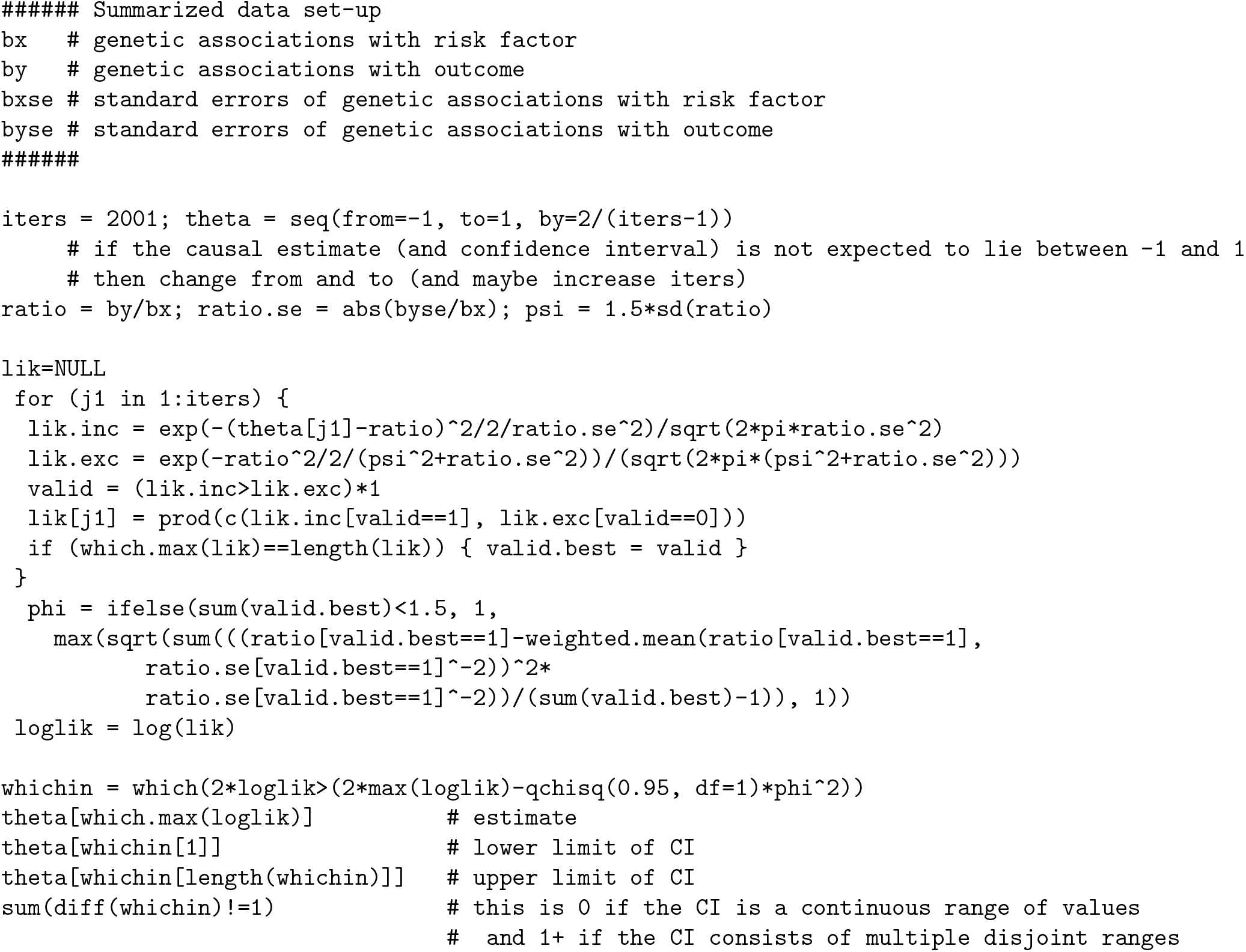

### A.2 Supplementary Figures and Tables

**Supplementary Figure A1:**
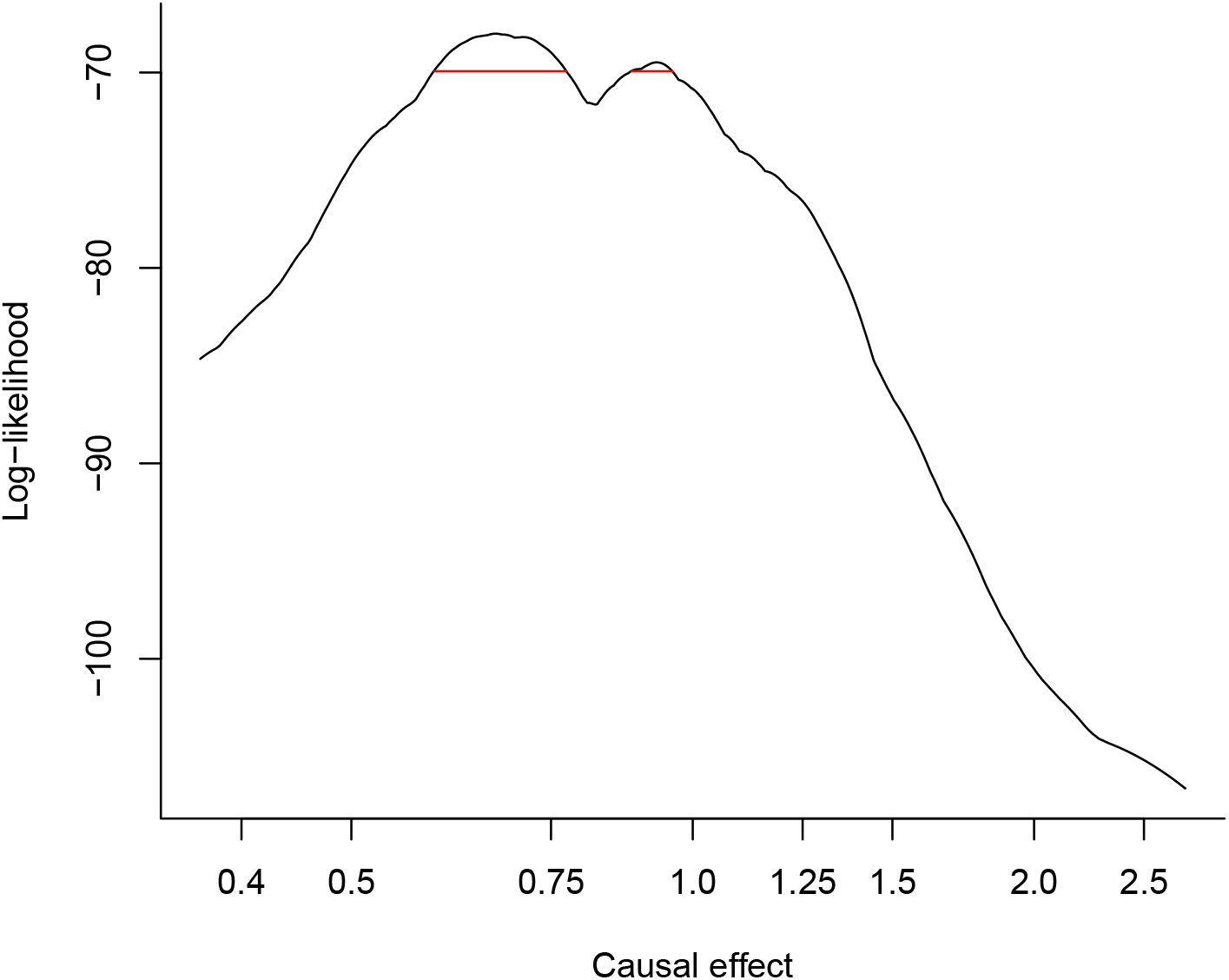
Log-likelihood function for applied example with recommended value of standard deviation 1.114. Red line indicates the 95% confidence interval for the causal effect. Causal estimate represents odds ratio for coronary heart disease per 1 standard deviation increase in HDL-cholesterol.

**Supplementary Figure A2:**
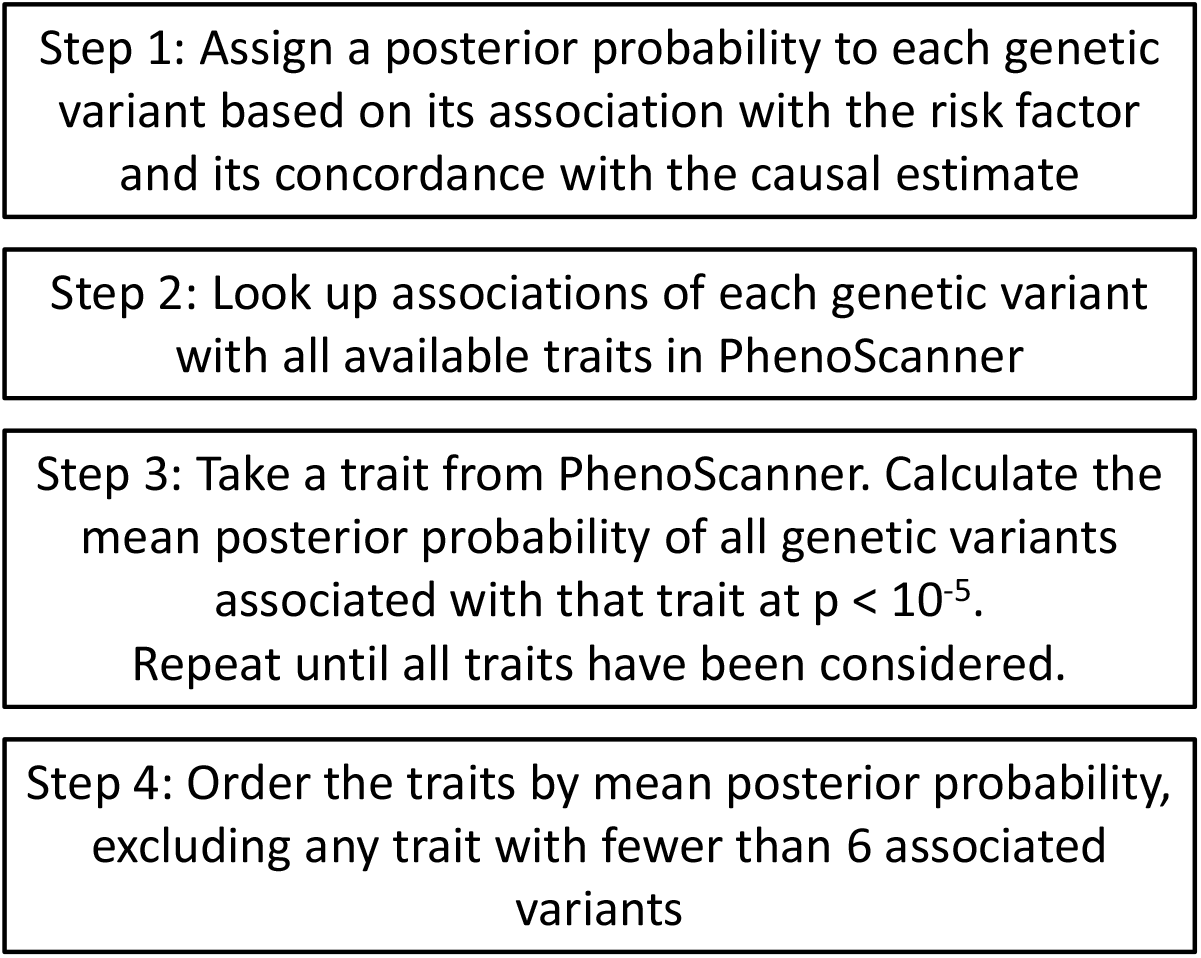
Flowchart illustrating approach for searching and ranking risk factors with available summarized data.

**Supplementary Figure A3:**
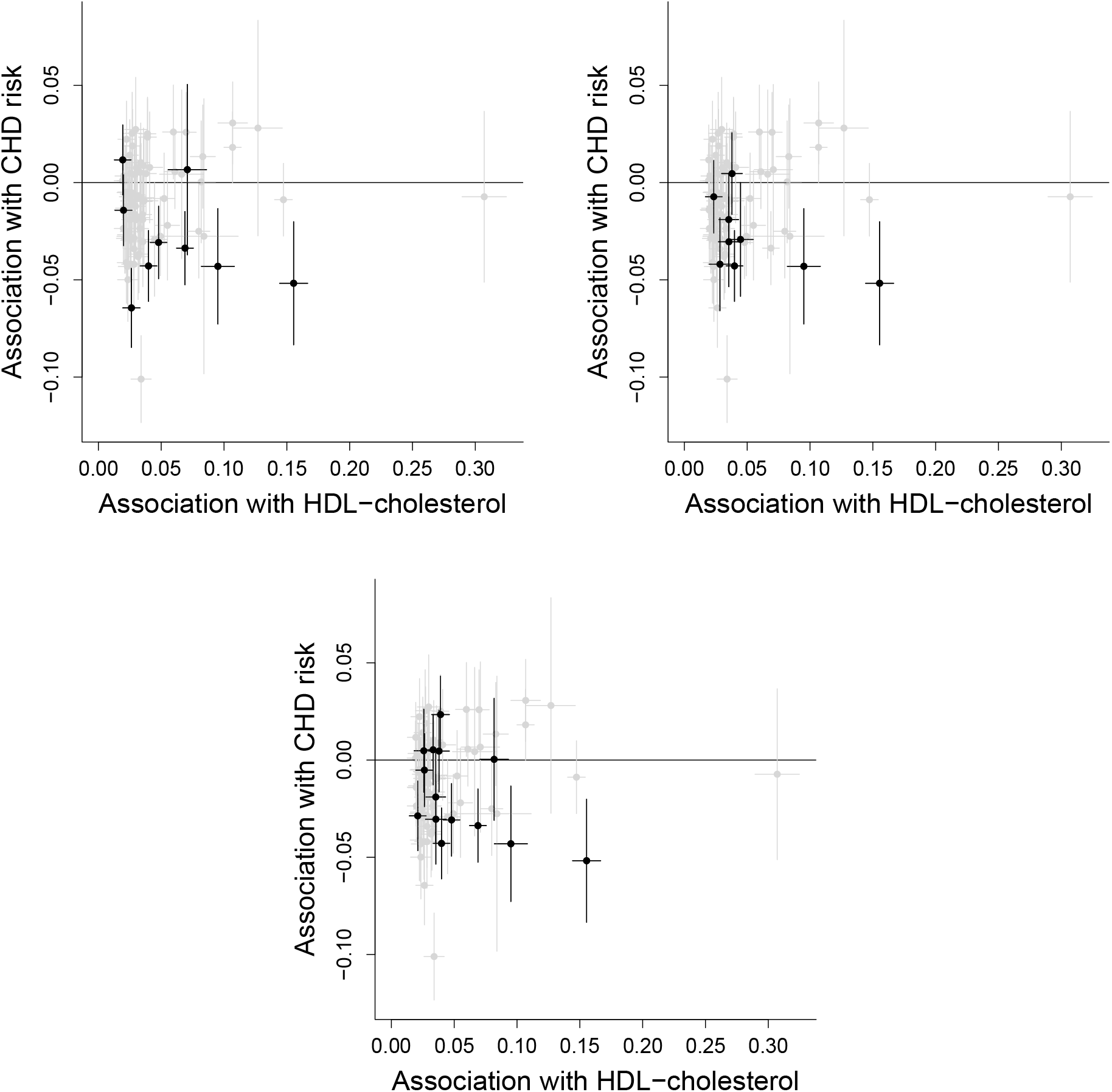
Genetic associations with HDL-cholesterol (standard deviation units) against genetic associations with CHD risk (log odds ratios). Variants in black are associated at a suggestive level of significance (p-value < 10^−5^) with: (top left) platelet distribution width, (top right) mean corpuscular hemoglobin concentration, and (bottom) red cell distribution width.

**Supplementary Table A1:**
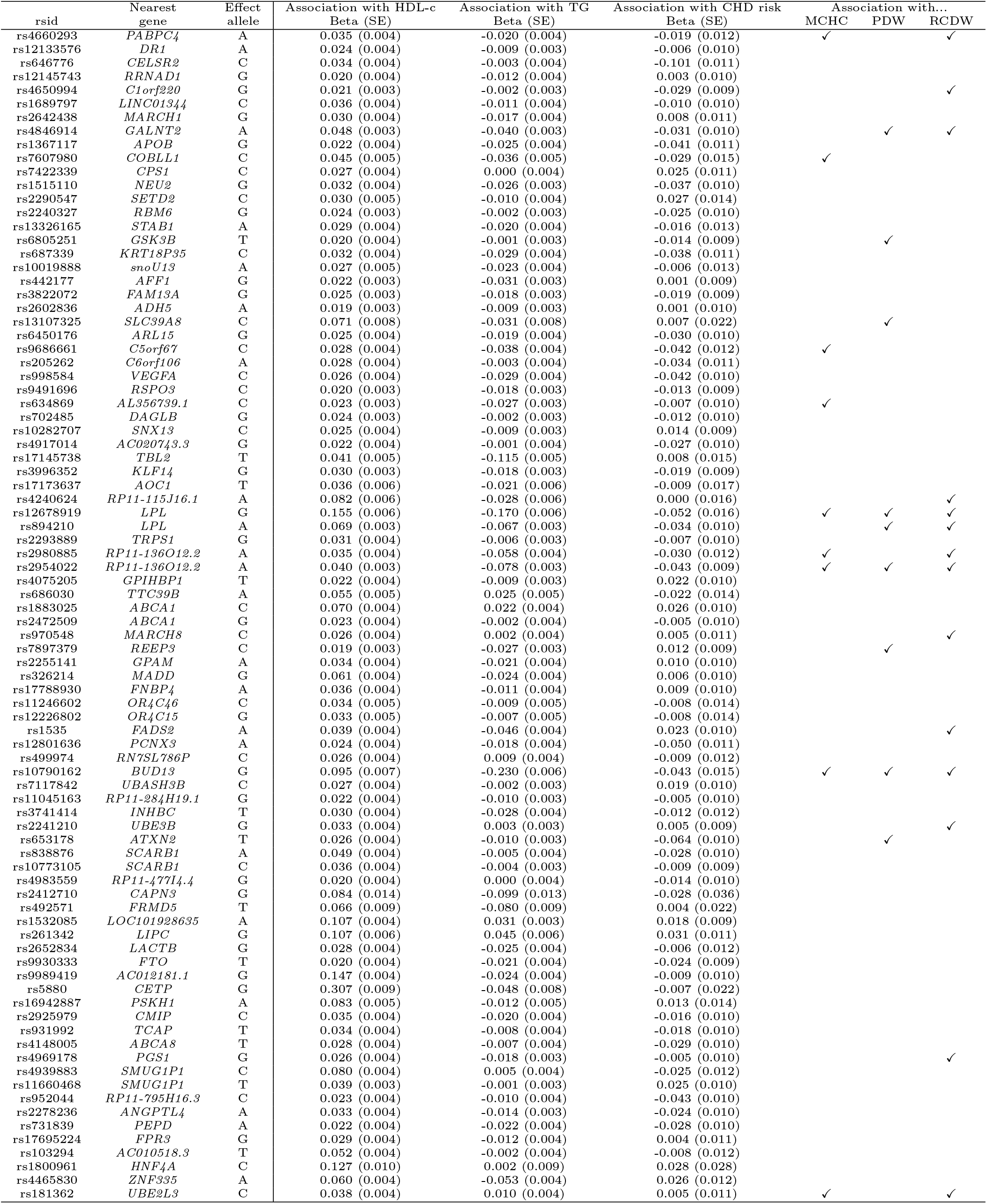
Details of genetic variants, beta-coefficients (standard errors, SE) for associations with HDL-cholesterol and triglycerides (TG) (both instandard deviation units) and with coronary heart disease (CHD) risk (log odds ratios), for 86 genetic variants. Tickmarks indicate associations at *p* < 10^−5^ with mean corpuscular hemoglobin concentration (MCHC), platelet distribution width (PDW), or red cell distribution width (RCDW).

**Supplementary Table A2:**
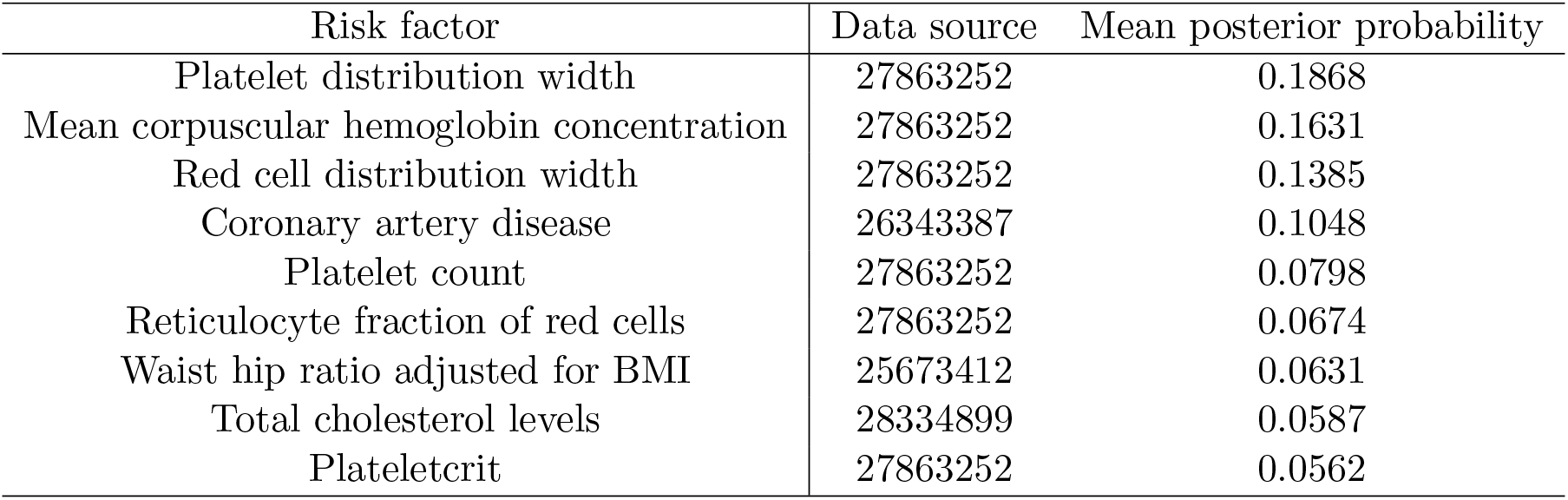
List of top risk factors from PhenoScanner ranked by mean posterior probability for applied example. Data source is the PubMed ID of the manuscript from which the association estimates were obtained.

**Supplementary Table A3:** Results from colocalization analysis using conservative and uniform settings for priors. Traits in parentheses only colocalized using uniform prior. For most gene regions, one set of traits that colocalized was identified; for *LPL* and *ATXN2*, two sets were identified. A checkmark indicates that colocalization was observed for the indicated choice of prior between HDL-cholesterol, CHD risk, and at least one of the blood cell traits. Abbreviations: HDL-c = HDL-cholesterol, TG = triglycerides, CHD = coronary heart disease risk, MCHC = mean corpuscular hemoglobin concentration, PDW = platelet distribution width, RCDW = red cell distribution width.

| Gene | Colocalization group 1 | Colocalization group 2 | Conservative | Uniform |
| --- | --- | --- | --- | --- |
| *C1orf220* | HDL-c, (CHD), MCHC, RCDW | - |  | ✓ |
| *GALNT2* | HDL-c, TG, (CHD), PDW, RCDW | - |  | ✓ |
| *COBLL1* | HDL-c, TG, MCHC, RCDW | - |  |  |
| *C5orf67* | HDL-c, TG, CHD, MCHC | - | ✓ | ✓ |
| *TRIB1* | HDL-c, TG, CHD, MCHC, PDW, RCDW | - | ✓ | ✓ |
| *LPL* | (HDL-c), CHD, PDW | TG, MCHC, RCDW |  | ✓ |
| *BUD13* | HDL-c, CHD, MCHC, PDW, RCDW | - | ✓ | ✓ |
| *ATXN2* | HDL-c, CHD | (MCHC), PDW, RCDW |  |  |
| *CMIP* | HDL-c, TG, (CHD), (MCHC) | - |  | ✓ |

**Supplementary Table A4:** Estimates and 95% confidence intervals from contamination mixture method for different values of the standard deviation parameter *ψ*. Estimates represent log odds ratios for coronary heart disease per 1 standard deviation increase in HDL-cholesterol. In some cases, the 95% confidence interval consists of two disjoint intervals.

| Standard deviation parameter | Estimate | 95% confidence interval |
| --- | --- | --- |
| $\psi = 0.8$ | -0.456 | -0.609, -0.332 |
| $\psi = 1.0$ | -0.422 | -0.535, -0.275 |
| $\psi = 1.2$ | -0.400 | -0.517, -0.251 and -0.154, -0.030 |
| $\psi = 1.5$ | -0.327 | -0.478, -0.245 and -0.193, -0.005 |
| $\psi = 1.8$ | -0.134 | -0.470, 0.003 |

## References

[1] Davey Smith G, Ebrahim S. ‘Mendelian randomization’: can genetic epidemiology contribute to understanding environmental determinants of disease? International Journal of Epidemiology 2003; 32(1):1–22, doi:10.1093/ije/dyg070.

[2] Burgess S, Thompson SG. Mendelian randomization: methods for using genetic variants in causal estimation. Chapman & Hall, Boca Raton, FL, 2015.

[3] Sanna S, van Zuydam NR, Mahajan A, Kurilshikov A, Vila AV, Võsa U, Mujagic Z, Masclee AA, Jonkers DM, Oosting M, et al. Causal relationships among the gut microbiome, short-chain fatty acids and metabolic diseases. Nature Genetics 2019; doi:10.1038/s41588-019-0350-x.

[4] Millard LA, Davies NM, Tilling K, Gaunt TR, Davey Smith G. Searching for the causal effects of body mass index in over 300 000 participants in UK Biobank, using Mendelian randomization. PLOS Genetics 2019; 15(2):e1007951, doi:10.1371/journal.pgen.1007951.

[5] Greenland S. An introduction to instrumental variables for epidemiologists. International Journal of Epidemiology 2000; 29(4):722–729, doi:10.1093/ije/29.4.722.

[6] Angrist J, Imbens G, Rubin D. Identification of causal effects using instrumental variables. Journal of the American Statistical Association 1996; 91(434):444–455, doi:10.2307/2291629.

[7] Greco M, Minelli C, Sheehan NA, Thompson JR. Detecting pleiotropy in Mendelian randomisation studies with summary data and a continuous outcome. Statistics in Medicine 2015; 34(21):2926–2940, doi:10.1002/sim.6522.

[8] Lawlor D, Harbord R, Sterne J, Timpson N, Davey Smith G. Mendelian randomization: using genes as instruments for making causal inferences in epidemiology. Statistics in Medicine 2008; 27(8):1133–1163, doi:10.1002/sim.3034.

[9] Davey Smith G, Ebrahim S. Mendelian randomization: prospects, potentials, and limitations. International Journal of Epidemiology 2004; 33(1):30–42, doi:10.1093/ije/dyh132.

[10] Nitsch D, Molokhia M, Smeeth L, DeStavola B, Whittaker J, Leon D. Limits to causal inference based on Mendelian randomization: a comparison with randomized controlled trials. American Journal of Epidemiology 2006; 163(5):397–403, doi:10.1093/aje/kwj062.

[11] VanderWeele T, Tchetgen Tchetgen E, Cornelis M, Kraft P. Methodological challenges in Mendelian randomization. Epidemiology 2014; 25(3):427–435, doi:10.1097/ede.0000000000000081.

[12] Kang H, Zhang A, Cai T, Small D. Instrumental variables estimation with some invalid instruments, and its application to Mendelian randomisation. Journal of the American Statistical Association 2016; 111(513):132–144, doi:10.1080/01621459.2014.994705.

[13] Windmeijer F, Farbmacher H, Davies N, Davey Smith G. On the use of the lasso for instrumental variables estimation with some invalid instruments. Technical Report Discussion Paper 16/674, University of Bristol 2016.

[14] Bowden J, Davey Smith G, Haycock PC, Burgess S. Consistent estimation in Mendelian randomization with some invalid instruments using a weighted median estimator. Genetic Epidemiology 2016; 40(4):304–314, doi:10.1002/gepi.21965.

[15] Guo Z, Kang H, Tony Cai T, Small DS. Confidence intervals for causal effects with invalid instruments by using two-stage hard thresholding with voting. Journal of the Royal Statistical Society: Series B (Statistical Methodology) 2018; 80(4):793–815.

[16] Hartwig FP, Davey Smith G, Bowden J. Robust inference in summary data Mendelian randomisation via the zero modal pleiotropy assumption. International Journal of Epidemiology 2017; 46(6):1985–1998, doi:10.1093/ije/dyx102.

[17] Verbanck M, Chen CY, Neale B, Do R. Detection of widespread horizontal pleiotropy in causal relationships inferred from Mendelian randomization between complex traits and diseases. Nature Genetics 2018; 50(5):693–698, doi:10.1038/s41588-018-0099-7.

[18] Bowden J, Davey Smith G, Burgess S. Mendelian randomization with invalid instruments: effect estimation and bias detection through Egger regression. International Journal of Epidemiology 2015; 44(2):512–525, doi:10.1093/ije/dyv080.

[19] Burgess S, Butterworth AS, Thompson SG. Mendelian randomization analysis with multiple genetic variants using summarized data. Genetic Epidemiology 2013; 37(7):658–665, doi:10.1002/gepi.21758.

[20] Wooldridge J. Introductory econometrics: A modern approach. Chapter 15: Instrumental variables estimation and two stage least squares. South-Western, Nashville, TN, 2009.

[21] Burgess S, Davies NM, Thompson SG. Bias due to participant overlap in two-sample Mendelian randomization. Genetic Epidemiology 2016; 40(7):597–608, doi:10.1002/gepi.21998.

[22] The global Lipids Genetics Consortium. Discovery and refinement of loci associated with lipid levels. Nature Genetics 2013; 45:1274–1283, doi:10.1038/ng.2797.

[23] CARDIoGRAMplusC4D Consortium. A comprehensive 1000 Genomes-based genome-wide association meta-analysis of coronary artery disease. Nature Genetics 2015; 47:1121–1130, doi:10.1038/ng.3396.

[24] Burgess S, Davey Smith G. Mendelian randomization implicates high-density lipoprotein cholesterol–associated mechanisms in etiology of age-related macular degeneration. Ophthalmology 2017; 124(8):1165–1174, doi:10.1016/j.ophtha.2017.03.042.

[25] Tardif JC, Rhéaume E, Lemieux Perreault LP, Grégoire JC, Feroz Zada Y, Asselin G, Provost S, Barhdadi A, Rhainds D, L’allier PL, et al. Pharmacogenomic determinants of the cardiovascular effects of dalcetrapib. Circulation: Cardiovascular Genetics 2015; 8(2):372–382, doi:10.1161/CIRCGENETICS.114.000663.

[26] Lincoff AM, Nicholls SJ, Riesmeyer JS, Barter PJ, Brewer HB, Fox KA, Gibson CM, Granger C, Menon V, Montalescot G, et al. Evacetrapib and cardiovascular outcomes in high-risk vascular disease. New England Journal of Medicine 2017; 376(20):1933–1942.

[27] HPS3/TIMI55–REVEAL Collaborative Group. Effects of anacetrapib in patients with atherosclerotic vascular disease. New England Journal of Medicine 2017; 377(13):1217–1227.

[28] Burgess S, Thompson SG. Multivariable Mendelian randomization: the use of pleiotropic genetic variants to estimate causal effects. American Journal of Epidemiology 2015; 181(4):251–260, doi:10.1093/aje/kwu283.

[29] Staley JR, Blackshaw J, Kamat MA, Ellis S, Surendran P, Sun BB, Paul DS, Freitag D, Burgess S, Danesh J, et al. PhenoScanner: a database of human genotype-phenotype associations. Bioinformatics 2016; 32(20):3207–3209, doi:10.1093/bioinformatics/btw373.

[30] van der Stoep M, Korporaal SJ, Van Eck M. High-density lipoprotein as a modulator of platelet and coagulation responses. Cardiovascular Research 2014; 103(3):362–371, doi:10.1093/cvr/cvu137.

[31] Burgess S, Zuber V, Gkatzionis A, Foley CN. Modal-based estimation via heterogeneity-penalized weighting: model averaging for consistent and efficient estimation in Mendelian randomization when a plurality of candidate instruments are valid. International Journal of Epidemiology 2018; 47(4):1242–1254, doi:10.1093/ije/dyy080.

[32] Stacey D, Fauman EB, Ziemek D, Sun BB, Harshfield EL, Wood AM, Butterworth AS, Suhre K, Paul DS. ProGeM: a framework for the prioritization of candidate causal genes at molecular quantitative trait loci. Nucleic Acids Research 2018; :gky837 doi:10.1093/nar/gky837.

[33] Didelez V, Sheehan N. Mendelian randomization as an instrumental variable approach to causal inference. Statistical Methods in Medical Research 2007; 16(4):309–330, doi:10.1177/0962280206077743.

[34] Thompson JR, Minelli C, Fabiola Del Greco M. Mendelian randomization using public data from genetic consortia. The International Journal of Biostatistics 2016; 12(2).

[35] Bowden J, Del Greco F, Minelli C, Lawlor D, Sheehan N, Thompson J, Davey Smith G. Improving the accuracy of two-sample summary data Mendelian randomization: moving beyond the NOME assumption. International Journal of Epidemiology 2018; Available online before print., doi:10.1093/ije/dyy258.

[36] Burgess S, Dudbridge F, Thompson SG. Combining information on multiple instrumental variables in Mendelian randomization: comparison of allele score and summarized data methods. Statistics in Medicine 2016; 35(11):1880–1906, doi:10.1002/sim.6835.

[37] Thompson S, Sharp S. Explaining heterogeneity in meta-analysis: a comparison of methods. Statistics in Medicine 1999; 18(20):2693–2708.

[38] Bowden J, Del Greco M F, Minelli C, Davey Smith G, Sheehan N, Thompson J. A framework for the investigation of pleiotropy in two-sample summary data Mendelian randomization. Statistics in Medicine 2017; 36(11):1783–1802, doi:10.1002/sim.7221.

[39] Giambartolomei C, Vukcevic D, Schadt EE, Franke L, Hingorani AD, Wallace C, Plagnol V. Bayesian test for colocalisation between pairs of genetic association studies using summary statistics. PLOS Genetics 2014; 10(5):e1004383, doi:10.1371/journal.pgen.1004383.

[40] Astle WJ, Elding H, Jiang T, Allen D, Ruklisa D, Mann AL, Mead D, Bouman H, Riveros-Mckay F, Kostadima MA, et al. The allelic landscape of human blood cell trait variation and links to common complex disease. Cell 2016; 167(5):1415–1429.

[41] Berisa T, Pickrell JK. Approximately independent linkage disequilibrium blocks in human populations. Bioinformatics 2016; 32(2):283–285, doi:10.1093/bioinformatics/btv546.

